# Fibroblasts impair muscle stem cell self-renewal via excessive fibronectin deposition in viscoelastic hydrogel co-cultures

**DOI:** 10.64898/2026.07.03.736419

**Authors:** Tze-Ling Chang, Tenaya K. Vallery, Thea S. Zlatkov, Bradley B. Olwin, Kristi S. Anseth

## Abstract

Muscle satellite cells (SCs) regenerate skeletal muscle, but their regenerative capacity declines with age, in part due to extracellular matrix (ECM) remodeling and aberrant fibroblast activation within the SC niche. In regenerating young mouse muscle, fibronectin remodeling is transient, whereas in aged mouse muscle, fibronectin remodeling is prolonged and disorganized. Fibroblasts in aged mice are activated, increasing fibronectin deposition and expressing elevated α-smooth muscle actin (αSMA), which negatively influence SC fate. We develop a viscoelastic hydrogel co-encapsulation system, enabling three-dimensional co-culture of intact myofibers with primary fibroblasts. Using this 3D co-culture system, we show that fibroblasts from young mice support SC quiescence and self-renewal, whereas fibroblasts from aged mice aberrantly activate SCs and promote their differentiation on myofibers isolated from either young or aged mice. Knocking down fibronectin (*Fn1*) in fibroblasts from aged mice partially restores SC function, promoting quiescence and limiting differentiation. Using a novel 3D hydrogel co-culture system, we demonstrate that fibroblast-deposited fibronectin is a key age-associated regulator negatively affecting SC fate within the SC niche of aged mice.

## 1. Introduction

Maintaining and regenerating skeletal muscle requires satellite cells (SCs), or muscle stem cells, which reside in a specialized niche between the basal lamina and the sarcolemma of muscle myofibers.^[1]^ Under homeostasis, SCs are typically quiescent; however, in response to muscle injury, SCs activate and re-enter the cell cycle. Activated SCs either self-renew to replenish the stem cell pool or proliferate to generate myogenic progenitors (myoblasts).^[2]^ Myoblasts differentiate and fuse to form new myofibers or repair damaged myofibers. During aging, muscle regeneration is compromised as SCs in aged organisms are reduced in number and impaired.^[3]^

While SC cell autonomous changes in aged organisms contribute to their functional decline,^[4-6]^ cell non-autonomous changes in the SC niche of aged mice further impair SC interactions with the surrounding microenvironment. Transplanting SCs from young mice into aged mice fails to rescue the age-associated SC self-renewal deficit,^[6]^ indicating cell non-autonomous changes in the niche impair muscle maintenance and repair. Fibronectin, a key extracellular matrix (ECM) glycoprotein, undergoes dynamic remodeling during aging and muscle regeneration.^[7, 8]^ Following injury, fibronectin gene expression is upregulated by 55-fold in SCs, leading to a 16-fold increase in fibronectin deposition in the SC niche, promoting symmetric SC expansion. In the SC niche of aged mice, reduced fibronectin levels impair SC function and maintenance;^[9]^ however, restoring fibronectin in the muscle of aged mice rescues SC activity and enhances SC regenerative capacity.^[9]^

Fibroblasts secrete fibronectin, playing a critical role in regenerating muscle following an injury.^[10, 11]^ Upon muscle damage, fibro-adipogenic progenitors (FAPs), a subset of fibroblasts, accumulate at the injury site and establish an inflammatory environment necessary for effective muscle repair.^[12]^ Ablating fibroblasts *in vivo* disrupts SC dynamics, leading to premature SC differentiation, SC pool depletion, and formation of smaller regenerated myofibers.^[13]^ Further, aging impairs FAPs, indirectly reducing SC myogenic potential;^[14]^ but how fibroblasts influence SC behavior and how aging affects SC-fibroblast interactions is poorly understood.

We employed a fully defined synthetic viscoelastic hydrogel to co-encapsulate fibroblasts and myofibers with their associated SCs to probe how fibroblasts influence SC behavior on myofibers and how aging alters this interaction. Our culture system recapitulates muscle-relevant viscoelastic properties, while preserving SC polarity and the native cell-myofiber-matrix interface.^[15]^ Using our culture system, we demonstrate that fibroblasts from aged mice are intrinsically activated, exhibiting elevated αSMA expression and increased fibronectin deposition, regardless of whether they are co-encapsulated with myofibers from young or aged mice. Moreover, fibroblasts from young mice enhance SC self-renewal, whereas fibroblasts from aged mice promote symmetric SC division and premature differentiation on co-encapsulated myofibers. Knockdown of *Fn1* in fibroblasts from aged mice partially rescues SC self-renewal and prevents SC from differentiating, suggesting that excessive fibronectin deposition in muscle from aged mice is a significant cell non-autonomous deficit contributing to SC aging phenotypes.

## 2. Results and Discussion

### 2.1. Transient fibronectin remodeling in young mouse muscle becomes prolonged and dysregulated with age

Fibronectin, an extracellular matrix (ECM) component, undergoes dynamic remodeling during muscle regeneration and critically regulates SC function.^[7, 8]^ Upon muscle injury, deposits of fibronectin increase up to 16-fold, promoting symmetric SC expansion.^[16]^ Reducing fibronectin in the SC niche of aged mice impairs SC function and maintenance.^[9]^ Despite these observations, the dynamic regulation of fibronectin deposition during regeneration in young and aged mouse muscle remain poorly understood.

Tibialis anterior (TA) muscles of wild-type young (4-6 months old) and aged (20-24 months old) mice were injured via BaCl_2_ injection and collected at 4 and 7 days post-injury (dpi), with uninjured controls. SCs were identified by Pax7 immunoreactivity and fibronectin deposition was visualized by immunoreactivity to fibronectin **(Figure 1A)**. In TA muscle from young mice, SC numbers increased from 101 ± 17 cells/mm^2^ before injury to 747 ± 46 cells/mm^2^ at 4 dpi and declined to 474 ± 97 cells/mm^2^ by 7 dpi, where SCs expand rapidly soon after injury, returning to uninjured numbers as regeneration is complete **(Figure 1B)**. In contrast, SC numbers in injured muscle from aged mice failed to expand to levels comparable to those observed in muscle of young mice at 4 dpi **(Figure 1B)**. In TA muscle from young mice, fibronectin coverage increased at 4 dpi (13.7 ± 1.2%) over that in uninjured muscle (6.1 ± 0.3%), but decreased by 7 dpi (9.0 ± 0.4%), identifying a transient fibronectin upregulation when SCs rapidly expand **(Figure 1C)**. In contrast, fibronectin coverage remained elevated in TA muscle of aged mice at 7 dpi (17.1 ± 2.9%), identifying prolonged fibronectin accumulation following injury. Fibronectin immunoreactivity was low in uninjured muscle from young mice (control), transiently increased following an injury, reaching a maximum at 4 dpi and then declining by 7 dpi (**Figure 1D**). In muscle from aged mice, fibronectin was elevated in uninjured tissue and remained elevated following an injury (**Figure 1D**).

**Figure 1.**
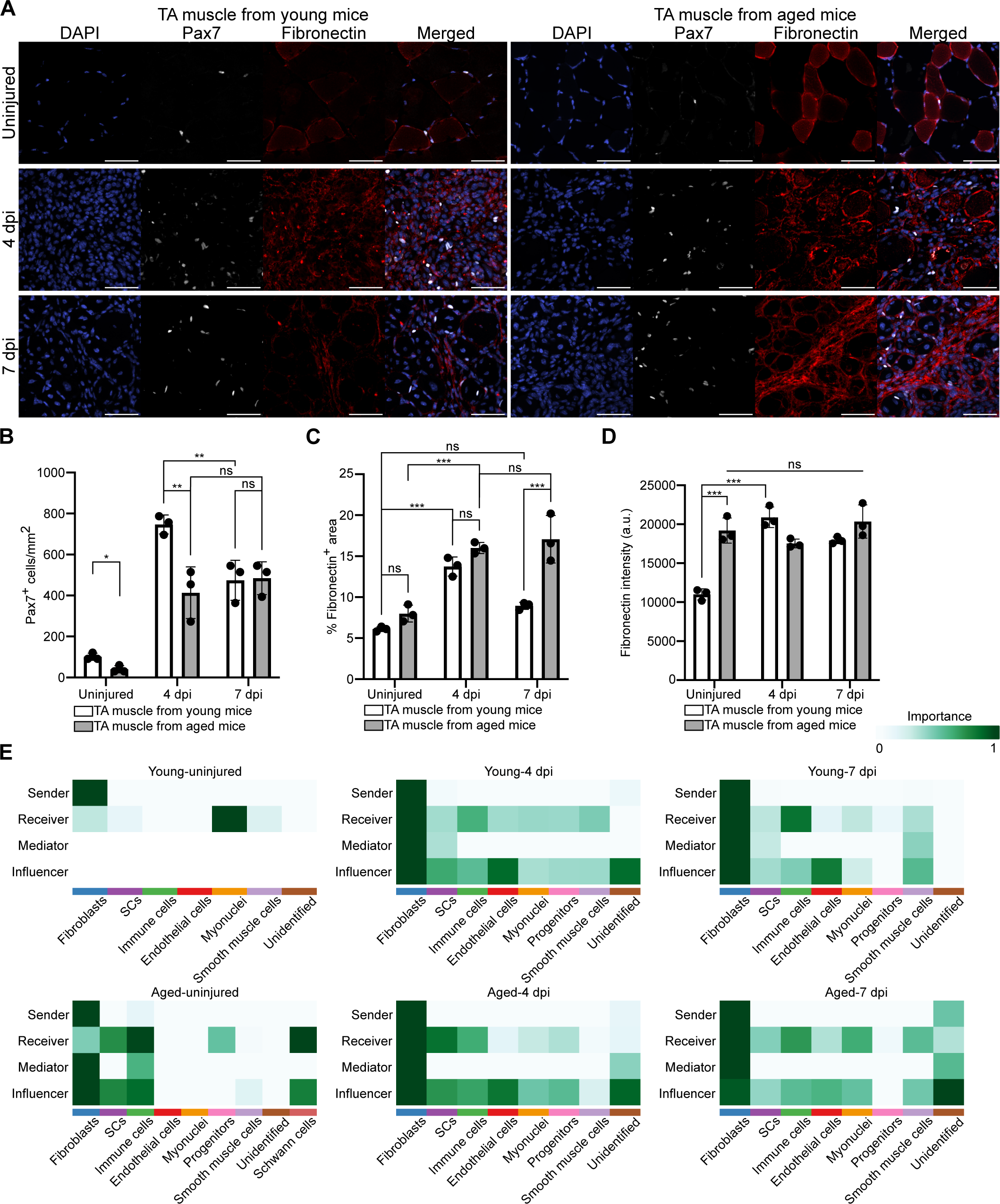
Fibronectin deposition and signaling transiently increases in muscle of young mice but persists in muscle of aged mice following an injury. (A) Representative images of tibialis anterior (TA) muscle sections from young and aged mice, either uninjured or at 4 and 7 days post-injury (dpi). Cell nuclei (DAPI, blue), SCs (Pax7 immunoreactivity, white), and fibronectin immunoreactivity (red). Scale bars, 100 μm. (B-D) SC numbers (B), fibronectin coverage (C), and fibronectin intensity (D) quantified in TA muscle sections from young and aged mice, either uninjured or at 4 and 7 dpi. *N* = 3 biological replicates; data are presented as mean ± SD; *\*P* < 0.05, *\*\*P* < 0.01, and *\*\*\*P* < 0.001 in a two-way ANOVA test. (E) *CellChat* network centrality analysis of fibronectin signaling pathways in young and aged mice in uninjured muscle, or at 4 and 7 dpi.

Fibronectin remodeling following an injury is dynamically regulated only in young mice, revealing distinct temporal patterns between muscles in young and aged mice.

Fibronectin remodeling appears transiently regulated in muscle of young mice, yet in aged mouse muscle, fibronectin deposition is dysregulated and persists. In young mouse muscle, transient increases in fibronectin after injury promote SC activation and proliferation,^[17]^ that decline by degrading the extracellular matrix (ECM) when muscle regeneration is complete, preventing muscle fibrosis.^[18]^ In contrast, in aged mouse muscle ECM remodeling is impaired where fibronectin accumulates and persists in the absence of an injury. Concomitant with fibronectin remodeling, additional hallmarks of chronic fibrosis are present in aged mouse skeletal muscle, including increased collagen deposition, elevated ECM stiffness, and age-related matrix cross-linking.^[19-21]^ A 3-fold increase in Young’s modulus was measured in TA muscles of aged mice compared to young counterparts.^[22]^ ECM derived from aged mice inhibits SC expansion, highlighting a critical role of the SC microenvironment in driving age-associated muscle phenotypes.^[23]^ The dysregulation of ECM turnover with age may reflect reduced matrix metalloproteinase activity or elevated TGF-β signaling, both of which promote fibroblast activation and fibrotic ECM deposition.^[24-26]^

Fibronectin immunoreactivity differed significantly between TA muscle from young and aged mice; therefore, we analyzed *CellChat* network centrality in previously published single-cell and single-nucleus RNA-sequencing datasets of TA muscle to identify cell types involved in fibronectin signaling (**Figure 1E**).^[27]^ *CellChat* infers intercellular communication based on established ligand-receptor interactions. Senders produce ligands to initiate signaling, receivers express the nominal receptors, mediators relay or amplify signals between senders and receivers, and influencers modulate the overall signaling dynamics. Fibroblasts are the dominant senders of fibronectin signaling across all cell type groups under all conditions. In uninjured muscle of aged mice, fibroblast signaling involvement was increased across all signaling roles compared to uninjured muscle of young mice. In addition, following injury, fibroblasts in muscles of young and aged mice upregulated fibronectin signaling and functioned predominantly as senders, receivers, mediators, and influencers, demonstrating that fibroblasts play a major role in fibronectin signaling during muscle repair.

### 2.2. Aged mouse muscle exhibits sustained accumulation of fibroblasts after injury

Since *CellChat* analysis identified fibroblasts as a predominant source of fibronectin signaling within the regenerative niche, we examined whether changes in fibroblast abundance and activation occur following injury in muscles from young and aged mice. TA muscles from wild-type young (4-6 months old) and aged (20-24 months old) mice were injured by BaCl_2_ injection, and tissues were collected at 4 and 7 dpi, as well as uninjured controls. Fibroblasts and activated fibroblasts were identified by platelet-derived growth factor receptor ⍺ (PDGFR⍺) and ⍺-smooth muscle actin (⍺SMA) immunoreactivity, respectively **(Figure 2A)**. In TA muscles of young mice, the number of PDGFR⍺^+^ fibroblasts increased at 4 dpi (60 ± 10 cells/mm^2^) compared to uninjured controls (290 ± 58 cells/mm^2^) and declined by 7 dpi (179 ± 35 cells/mm^2^) **(Figure 2B)**. In TA muscles of aged mice, PDGFR⍺^+^ fibroblast numbers increased at 4 dpi (180 ± 32 cells/mm^2^) and further increased by 7 dpi (278 ± 11 cells/mm^2^), with a similar trend observed for ⍺SMA^+^ activated fibroblasts **(Figure 2C)**. Greater numbers of PDGFR⍺^+^ fibroblasts and ⍺SMA^+^ activated fibroblasts were present in uninjured TA muscles from aged mice than in uninjured TA muscles from young mice. Together, fibroblasts and activated fibroblasts in young mouse muscle are transiently enriched at the injury site, peaking at 4 dpi and resolving by 7 dpi, whereas in aged mouse muscle, fibroblast accumulation is prolonged and remains elevated throughout the 7 days following injury.

**Figure 2.**
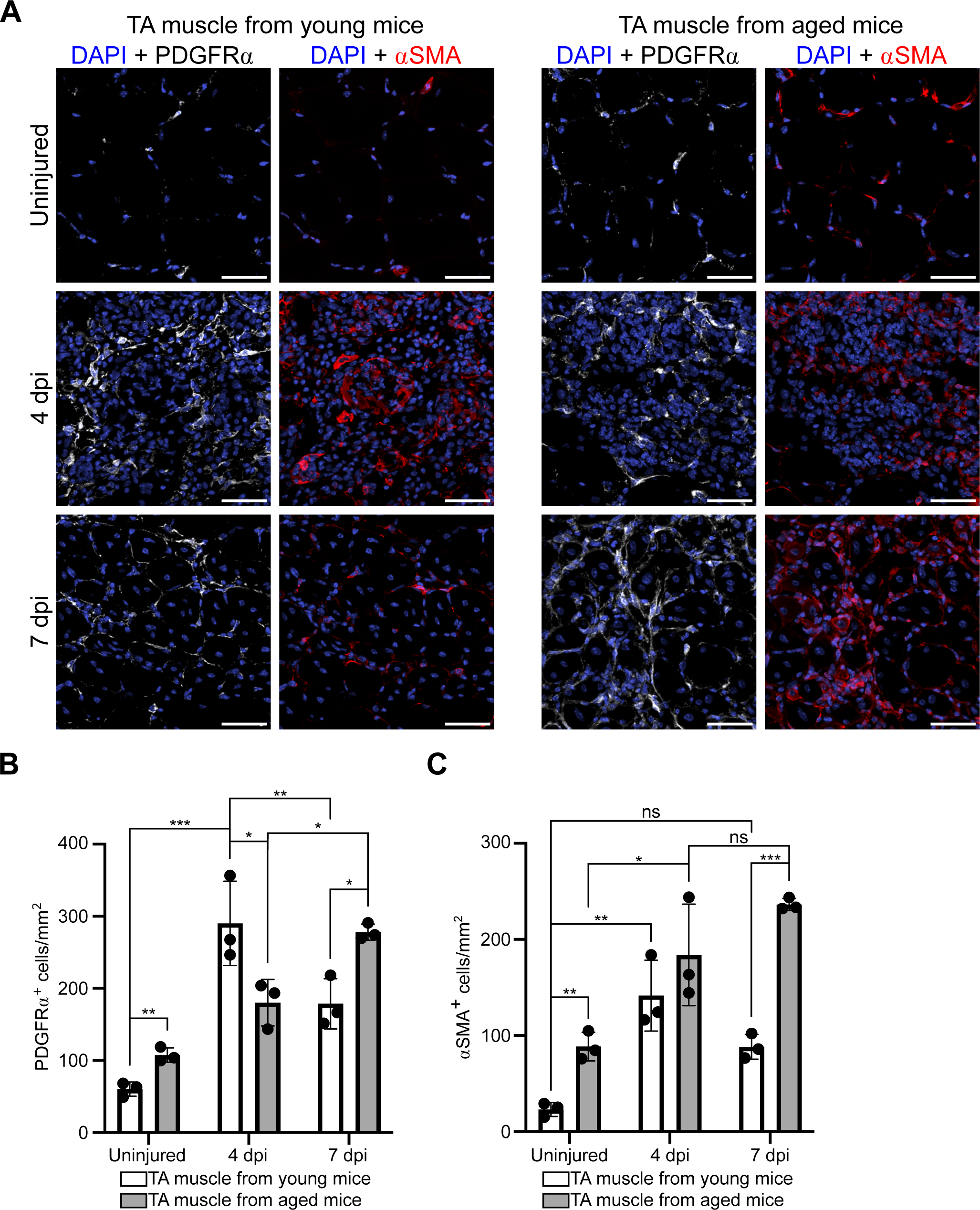
Fibroblast accumulation is transient in muscle of young mice but prolonged in aged mouse muscle. (A) Representative images of TA muscle sections from young and aged mice, either uninjured or at 4 and 7 dpi. Cell nuclei identified by DAPI staining (blue), fibroblasts by PDGFR⍺ immunoreactivity (white), and activated fibroblasts by αSMA immunoreactivity (red). Scale bars, 100 μm. (B-C) Fibroblast numbers (B) and activated fibroblast numbers (C) quantified in TA muscle sections from young and aged mice, either uninjured or at 4 and 7 dpi. *N* = 3 biological replicates; data are presented as mean ± SD; *\*P* < 0.05, *\*\*P* < 0.01, and *\*\*\*P* < 0.001 in a two-way ANOVA test.

### 2.3. Fibroblasts from aged mice are intrinsically activated and deposit increased fibronectin

After establishing differences in fibronectin deposition and fibroblast abundance between regenerating muscles of young and aged mice, we developed an *in vitro* system to delineate fibroblast-specific effects from the *in vivo* environment and assess how fibroblast contributions change during aging. Given that we observed a higher proportion of activated fibroblasts in TA muscles of aged mice, we first asked whether age-associated fibroblast phenotypes could be observed *in vitro*.

Fibroblasts isolated from young and aged mice were cultured on gelatin-coated plates for 2 days and assessed for αSMA and fibronectin immunoreactivity **(Figure 3A)**. Gelatin-coated tissue culture plates were used for 2D culture as a standard substrate that supports fibroblast adhesion through integrin-binding motifs similar to collagen without the added signaling bias of purified ECM proteins.^[28]^ Fibroblasts derived from aged mice markedly elevated their ⍺SMA levels compared to fibroblasts from young mice **(Figure 3B)**. αSMA⁺ fibroblasts, commonly referred to as myofibroblasts, are activated and associated with wound healing and fibrotic remodeling.^[29, 30]^ The proportion of ⍺SMA^+^ fibroblasts increased from 26.3 ± 4.7% in cultures of fibroblasts from young mice to 74.9 ± 3.6% in those from aged mice **(Figure 3C)**, demonstrating that fibroblasts from aged mice are more activated than those from young mice *in vitro*. Further, elevated fibronectin intensity was observed in cultures of fibroblasts from aged mice compared to those from young mice, likely related to the increased secretory properties of the ⍺SMA^+^ fibroblasts **(Figure 3D)**. Deposited fibronectin coverage increased from 3.0 ± 0.2% in cultures of fibroblasts from young mice to 10.2 ± 1.4% in those from aged mice **(Figure 3E)**.

**Figure 3.**
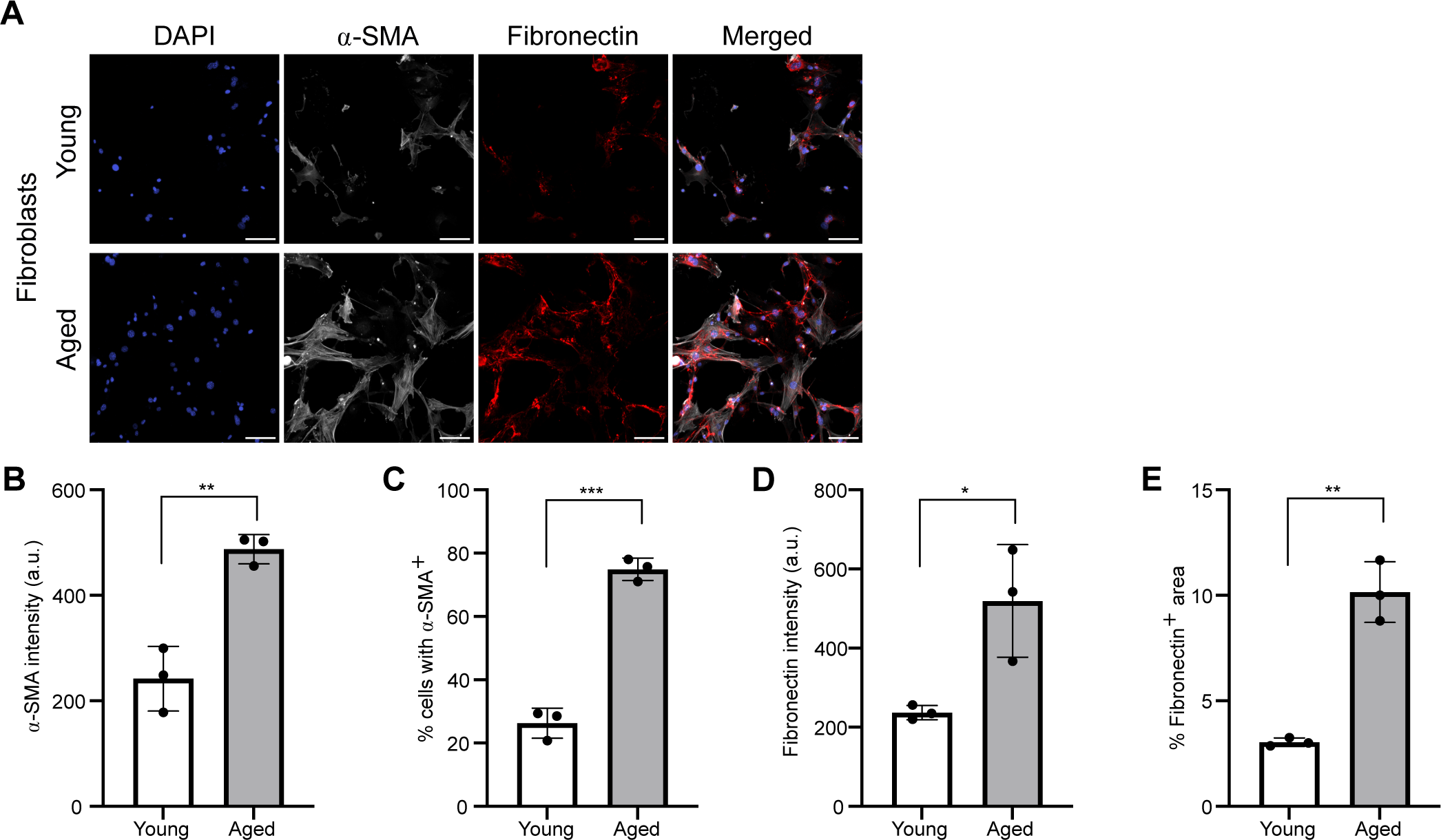
Fibroblasts isolated from aged mice are intrinsically activated *in vitro*. (A) Representative images of fibroblasts from young and aged mice cultured on gelatin-coated plates and assessed for ⍺SMA and fibronectin immunoreactivity after 2 days. Scale bars, 100 μm. (B-E) Quantification of ⍺SMA immunoreactivity (B), the percentage of ⍺SMA^+^ cells (C), fibronectin immunoreactivity (D), and the percentage of fibronectin^+^ area (E) in gelatin-coated plate cultures of fibroblasts from young or aged mice. *N* = 3 biological replicates; data are presented as mean ± SD; *\*P* < 0.05, *\*\*P* < 0.01, and *\*\*\*P* < 0.001 in a two-tailed unpaired student’s *t*-test.

Fibroblasts from aged mice appear intrinsically activated, as indicated by elevated ⍺SMA expression and increased fibronectin production, *in vivo* as well as *in vitro*. The persistently activated fibroblasts in aged mouse muscle resemble fibroblast behavior in fibrotic diseases including liver cirrhosis and pulmonary fibrosis, where myofibroblasts maintain elevated secretory activity and drive chronic wound healing responses, excessive ECM deposition, and tissue stiffening.^[31, 32]^ Similarly, in solid tumors including breast and pancreatic cancers, cancer-associated fibroblasts remain persistently activated, promoting stromal remodeling, fibrosis, and tumor progression.^[33, 34]^ Therefore, fibroblasts in muscles of aged mice adopt a persistent myofibroblast phenotype, as described in fibrotic tissues,^[35]^ likely leading to aberrant ECM remodeling that impairs SC function and muscle regeneration.

### 2.4. A fully defined viscoelastic hydrogel supports myofiber-fibroblast co-encapsulation

To decouple fibroblast-specific effects from other cell types in the *in vivo* environment, we established an *in vitro* myofiber-fibroblast co-encapsulation model. Mechanical properties of native muscle tissue are dynamic and essential for maintaining SC function and regulating cellular responses;^[15, 36, 37]^ unsurprisingly, culturing freshly isolated SCs leads to their spontaneous activation and differentiation due to the absence of matrix interactions. To address this limitation, we employed a hydrogel formulation, previously optimized for myofiber culture, that preserves SC quiescence and maintains SC-myofiber-matrix interactions.^[15]^

Specifically, hyaluronic acid (HA) was functionalized with hydrazide (R_1_) or alkyl-aldehyde (R_3_) groups, while a small fraction of HA-hydrazide pendant groups was further modified with azides (R_2_) **(Figure 4A and 4B)**.^[38]^ Hydrazide and alkyl-aldehyde groups react to form adaptable hydrazone bonds, imparting viscoelastic properties, while the azide groups react with 8-arm PEG-bicyclononyne (R_4_) via a strain-promoted azide-alkyne cycloaddition reaction to introduce permanent crosslinks into the network **(Figure 4C)**. We further introduced RGD peptides as pendant functional groups; RGD is an adhesive ligand derived from fibronectin and supports cell-matrix interactions.

**Figure 4.**
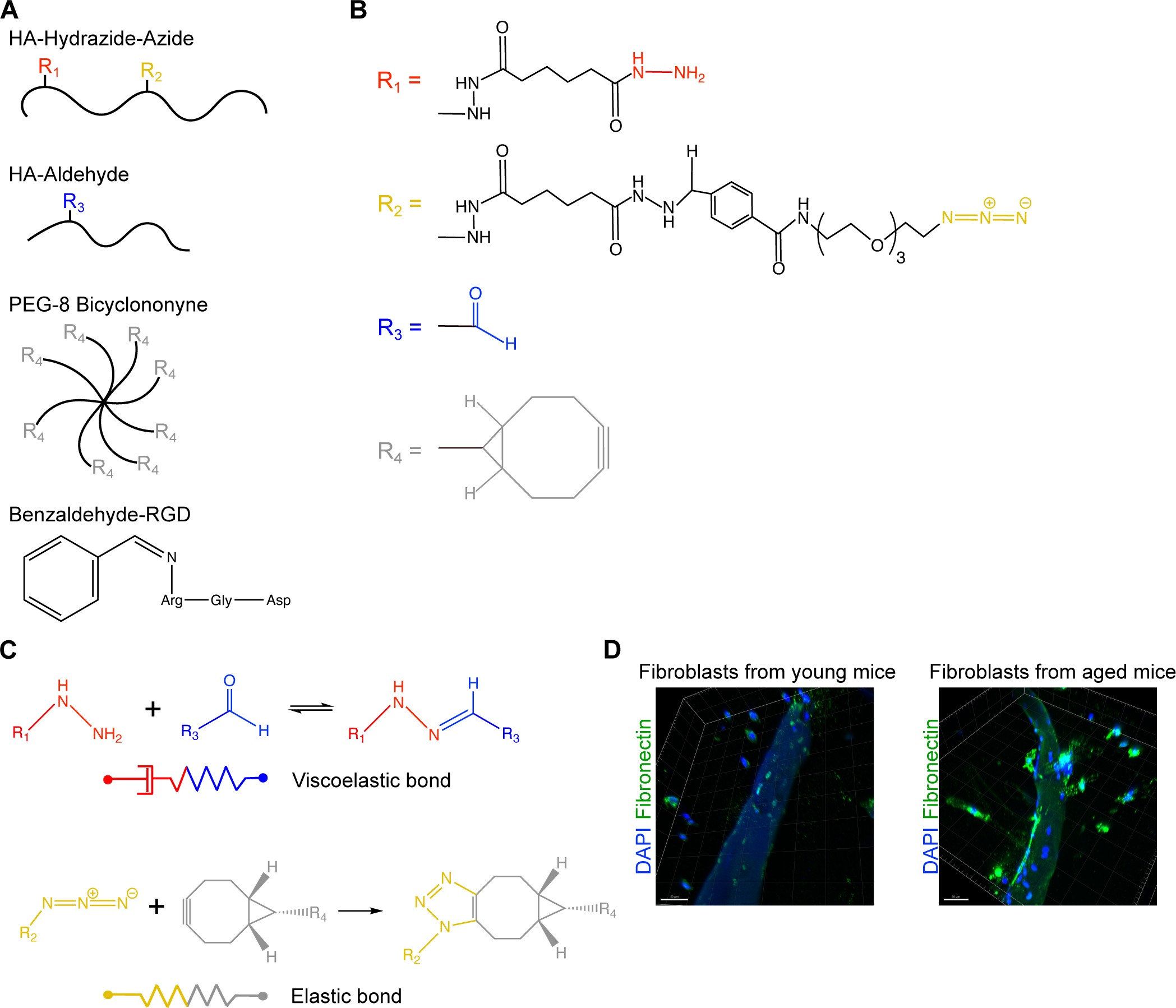
Network structure of the hydrogel for co-encapsulating fibroblasts and myofibers. (A) Schematic structures of 86 kDa HA-hydrazide-azide, 148 kDa HA-aldehyde, 40 kDa 8-arm PEG-BCN, and benzaldehyde-functionalized RGD ligand. (B and C) Chemical structures of functional groups (B) and crosslinks (C), including the alkyl-hydrazone bond (viscoelastic bond) and the triazole bond (elastic bond) in the hydrogel. (D) Representative images of fibroblasts from young or aged mice, co-encapsulated with myofibers in hydrogels with 1 mM RGD, labeled with fibronectin after 2 days in culture. Scale bars, 50 μm.

The final hydrogel formulation consisted of 5 wt% total polymer content with 88% adaptable hydrazone crosslinks. These viscoelastic hydrogels had a swollen shear modulus of 800 ± 110 Pa and relaxed 57 ± 6% of an applied stress over 600 seconds (**Table 1)**. Additional mechanical properties are summarized in **Table 1**.

**Table 1.**
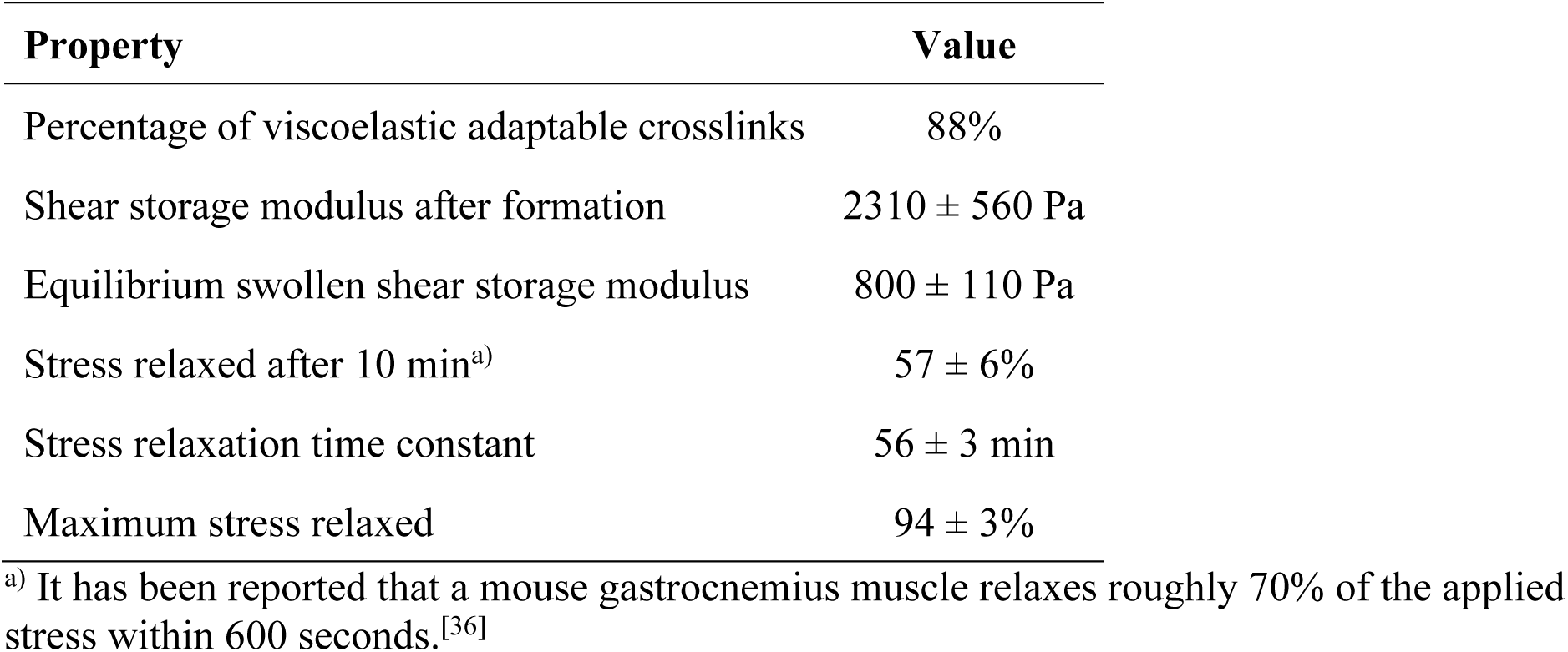
Mechanical properties of the hydrogel.

| Property | Value |
| --- | --- |
| Percentage of viscoelastic adaptable crosslinks | 88% |
| Shear storage modulus after formation | $2310 \pm 560$ Pa |
| Equilibrium swollen shear storage modulus | $800 \pm 110$ Pa |
| Stress relaxed after 10 min<sup>a)</sup> | $57 \pm 6\%$ |
| Stress relaxation time constant | $56 \pm 3$ min |
| Maximum stress relaxed | $94 \pm 3\%$ |

<sup>a)</sup> It has been reported that a mouse gastrocnemius muscle relaxes roughly 70% of the applied stress within 600 seconds.<sup>[36]</sup>

Fibroblasts isolated from young and aged mice were co-encapsulated with myofibers from young mice in hydrogels containing 1 mM RGD, cultured for 2 days, and then fixed and assessed for fibronectin immunoreactivity **(Figure 4D)**. Fibroblasts from aged mice produced substantially higher levels of fibronectin that accumulated along the myofiber surface and within pericellular regions. In contrast, fibronectin organization in cultures with fibroblasts isolated from young mice was diffuse and exhibited low immunoreactivity. Thus, the differences between muscle fibroblasts from young and aged mice observed *in vivo* and in 2D culture are retained within 3D hydrogel microenvironments, where fibroblasts isolated from aged mice maintain an intrinsically activated, matrix-producing phenotype. The engineered viscoelastic hydrogel system thus provides a physiologically relevant platform to test hypotheses and interrogate the effects of fibroblast-driven ECM remodeling and age-associated changes as muscle regenerates.

### 2.5. Fibroblasts from aged mice deposit more fibronectin regardless of the age of the myofiber donor

We co-encapsulated fibroblasts with myofibers using synthetic viscoelastic hydrogels to investigate whether the age of the myofiber donor influences fibroblast activation and fibronectin deposition. Fibroblasts from young and aged mice were co-encapsulated with myofibers isolated from either young or aged mice within hydrogels containing 1 mM RGD. These constructs were cultured for 2 days, then fixed and assayed for fibronectin and ⍺SMA immunoreactivity **(Figure 5A)**. Markedly elevated ⍺SMA expression was observed in fibroblasts from aged mice compared to those from young mice, regardless of the age of the myofiber donor **(Figure 5B)**. The proportion of ⍺SMA^+^ cells increased significantly from 6.0 ± 1.4% and 8.4 ± 3.3% in fibroblasts from young mice co-cultured with myofibers from young and aged mice, respectively, to 65.6 ± 8.6% and 70.7 ± 3.9% in fibroblasts from aged mice co-cultured with myofibers from young and aged mice, respectively **(Figure 5C)**.

**Figure 5.**
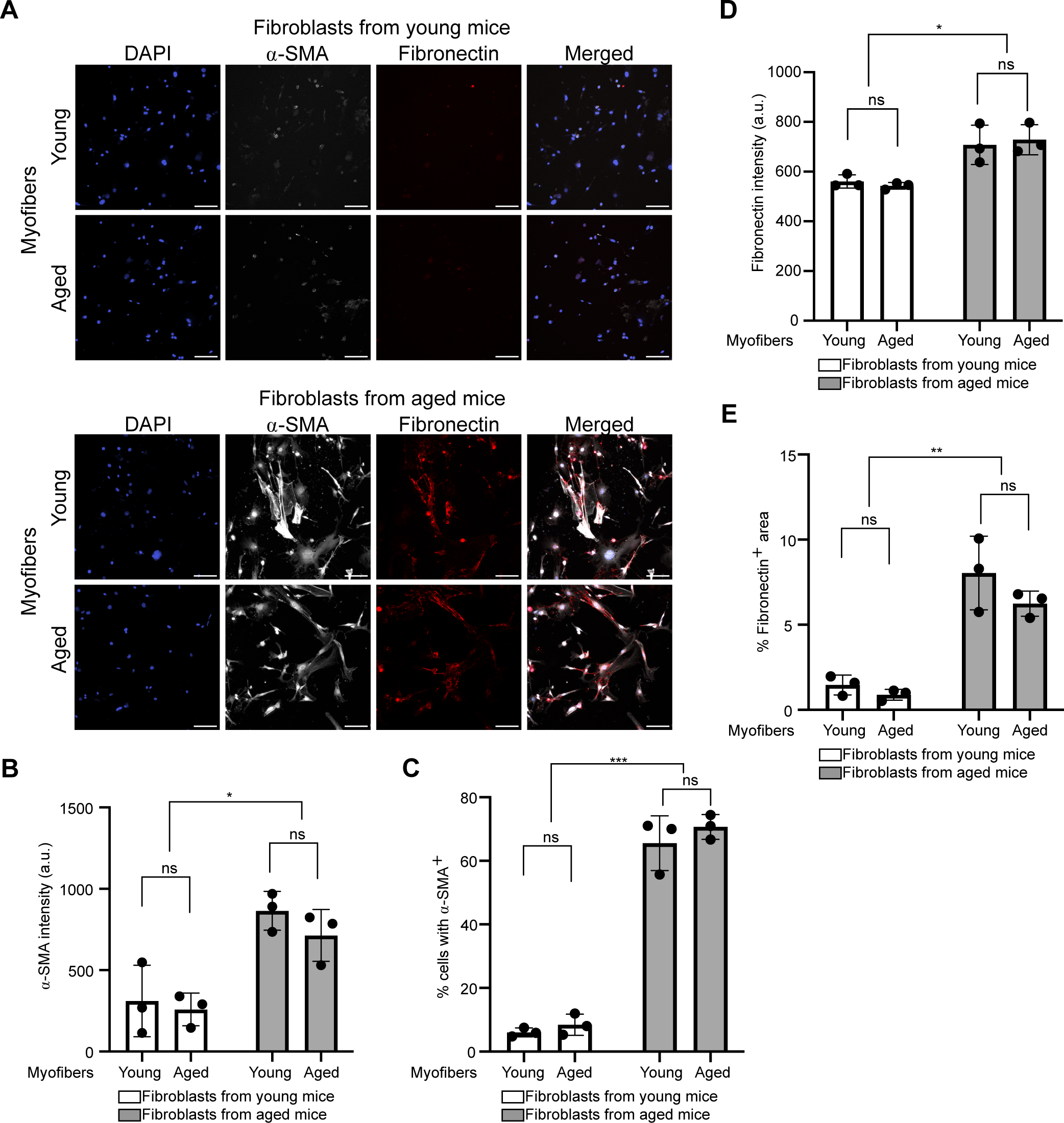
⍺SMA expression and fibronectin deposition are higher in fibroblasts from aged mice, regardless of co-encapsulation with myofibers from young or aged mice. (A) Representative images of fibroblasts from young or aged mice, co-encapsulated with myofibers from young or aged mice in hydrogels with 1 mM RGD, labeled with ⍺SMA and fibronectin after 2 days in culture. Scale bars, 100 μm. (B-E) Quantification of ⍺SMA immunoreactivity (B), the percentage of ⍺SMA^+^ cells (C), fibronectin immunoreactivity (D), and the percentage of fibronectin^+^ area (E) in fibroblasts from young or aged mice co-encapsulated with myofibers from young or aged mice in hydrogels with 1 mM RGD. *N* = 3 biological replicates; data are presented as mean ± SD; *\*P* < 0.05, *\*\*P* < 0.01, and *\*\*\*P* < 0.001 in a two-way ANOVA test.

Fibroblast activation appears primarily dependent on the age of the fibroblast donor rather than the age of the myofiber donor. Furthermore, fibronectin deposition by fibroblasts from aged mice was significantly higher than that of fibroblasts from young mice, as indicated by fibronectin intensity **(Figure 5D)** and fibronectin coverage area **(Figure 5E)**. Consistent with our *in vivo* observations, fibroblasts from aged mice maintain an intrinsically activated, fibronectin-rich phenotype, independent of whether they are cultured with myofibers from young or aged mice. Based on these findings, we next tested whether fibroblast-intrinsic aging phenotypes are sufficient to alter SC fate cultured on intact myofibers.

### 2.6. Fibroblasts from young mice promote SC self-renewal whereas fibroblasts from aged mice induce SC differentiation

We asked if fibroblasts from young or aged mice would influence SC fate decisions on myofibers from young and aged mice by co-encapsulating them within viscoelastic hydrogels containing 1 mM RGD. These constructs were cultured for two days, with a 2-hour EdU pulse administered prior to fixation to quantify proliferating SCs and determine cell state by Pax7 (stemness) and MyoD (activation) immunoreactivity **(Figure 6A and 6B)**.

**Figure 6.**
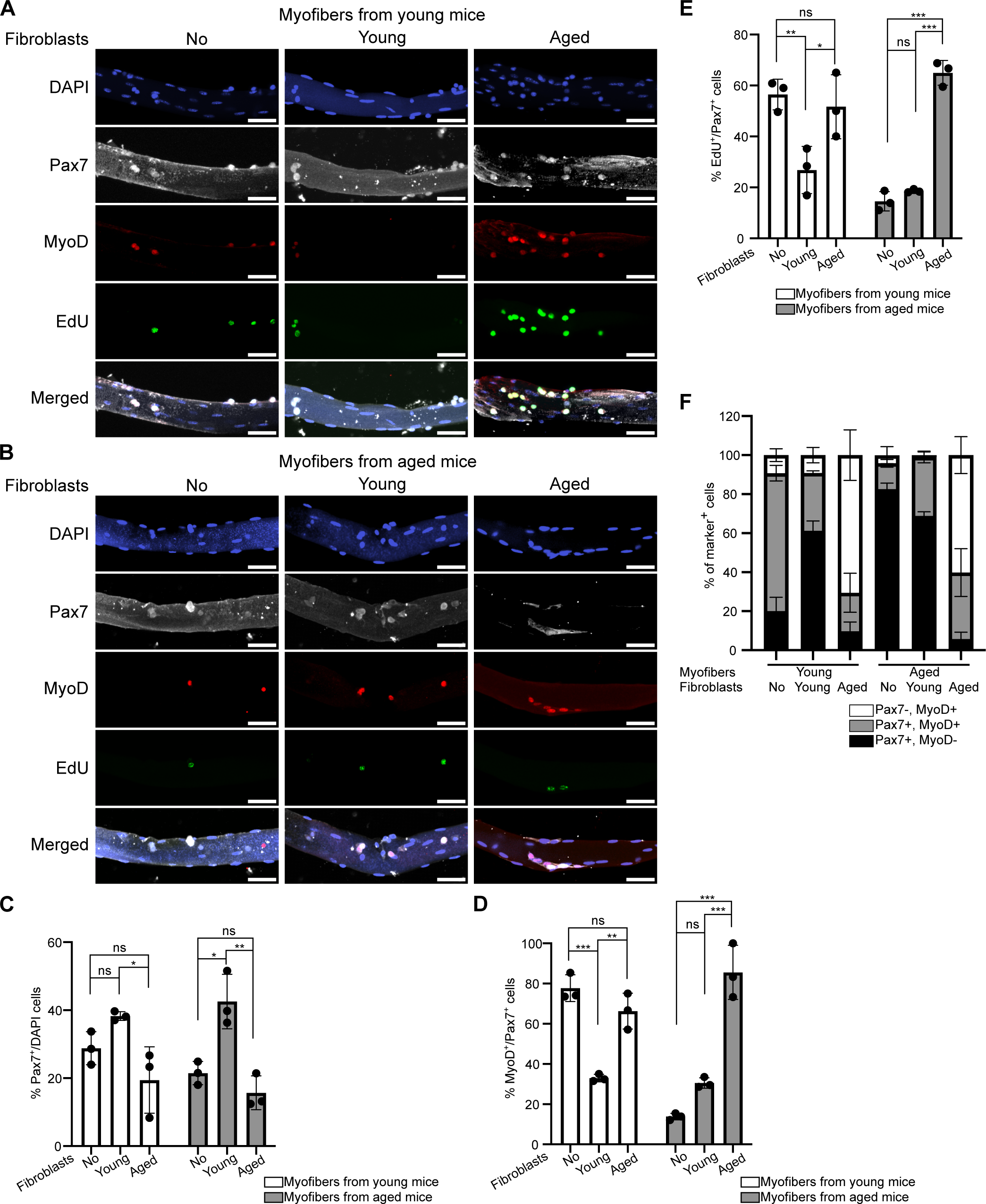
Fibroblasts from young mice enhance SC self-renewal, while fibroblasts from aged mice promote SC symmetric expansion. (A) Representative images of young mouse myofibers encapsulated in hydrogels with 1 mM RGD alone or co-encapsulated with fibroblasts from young or aged mice, SCs identified by Pax7 and MyoD immunoreactivity, and assessed for EdU incorporation after 2 days in culture. Scale bars, 50 μm. (B) Representative images of aged mouse myofibers encapsulated in hydrogels with 1 mM RGD alone or co-encapsulated with fibroblasts from young or aged mice, SCs identified by Pax7 and MyoD immunoreactivity, and assessed for EdU incorporation after 2 days in culture. Scale bars, 50 μm. (C-E) Quantification of Pax7^+^ (C), MyoD^+^ (D), and EdU^+^ (E) SCs on myofibers from young or aged mice, cultured alone or co-encapsulated with fibroblasts from young or aged mice in hydrogels with 1 mM RGD. *n* > 9 myofibers analyzed with samples from 3 mice, and statistics performed based on 3 independent mice; data are presented as mean ± SD; *\*P* < 0.05, *\*\*P* < 0.01, and *\*\*\*P* < 0.001 in a two-way ANOVA test. (F) Percentage of Pax7+/MyoD–, Pax7+/MyoD+, and Pax7–/MyoD+ cells on myofibers from young or aged mice, cultured alone or co-encapsulated with fibroblasts from young or aged mice in hydrogels with 1 mM RGD. *n* > 9 myofibers analyzed with samples from 3 mice, and data are presented as mean ± SD.

On myofibers from young mice, > 25% of DAPI^+^ nuclei were Pax7⁺ when encapsulated in hydrogels alone or with fibroblasts from young mice, decreasing to 19 ± 10% when co-encapsulated with fibroblasts from aged mice **(Figure 6C)**. On myofibers from aged mice, the percentage of Pax7⁺ cells increased from 21 ± 3% when encapsulated alone to 43 ± 8% with fibroblasts from young mice, but decreased dramatically to 16 ± 5% with fibroblasts from aged mice. Among Pax7^+^ cells on myofibers from young mice, 78 ± 7% were MyoD^+^ when encapsulated alone, decreasing markedly to 33 ± 2% with fibroblasts from young mice, while remaining relatively high at 66 ± 9% with fibroblasts from aged mice **(Figure 6D)**. On myofibers from aged mice, the percentage of MyoD⁺ cells increased from 14 ± 2% when encapsulated alone to 31 ± 3% with fibroblasts from young mice, but this increased over 5-fold to 86 ± 13% with fibroblasts from aged mice. More than half (56 ± 6%) of Pax7^+^ cells on myofibers from young mice were EdU^+^ when encapsulated alone, decreasing markedly to 27 ± 9% EdU^+^ with fibroblasts from young mice, but not changing significantly (52 ± 13% EdU^+^) with fibroblasts from aged mice **(Figure 6E)**. On myofibers from aged mice, fewer than 20% of Pax7⁺ cells were EdU⁺ when encapsulated alone or with fibroblasts from young mice, but this percentage increased to 65 ± 5% when co-encapsulated with fibroblasts from aged mice.

A minority (20 ± 7%) of SCs on myofibers from young mice were quiescent (Pax7+/MyoD–), 71 ± 4% were myoblasts (Pax7+/MyoD+), and 9 ± 3% were committed to terminal differentiation (Pax7–/MyoD+) when encapsulated alone **(Figure 6F)**. When co-encapsulated with fibroblasts from young mice, the proportion of quiescent SCs increased to 61 ± 5%, while myoblasts decreased to 30 ± 1%. In contrast, co-encapsulation with fibroblasts from aged mice led to a much lower percentage of quiescent SCs (10 ± 5%) and myoblasts (20 ± 10%), which was accompanied by a substantial 8-fold increase in differentiated myoblasts (71 ± 13%). On myofibers from aged mice, ∼75% of the SCs remained quiescent and ∼25% were either myoblasts or differentiated myoblasts when encapsulated alone or co-encapsulated with fibroblasts from young mice. However, co-encapsulation with fibroblasts from aged mice dramatically reduced the proportion of quiescent SCs to 6 ± 4% and increased differentiated myoblasts to 60 ± 9% on myofibers from aged mice.

Co-encapsulation of myofibers with fibroblasts from young mice increases the number of Pax7⁺ cells while limiting activation and proliferation, thereby promoting SC self-renewal. In contrast, co-encapsulation with fibroblasts from aged mice enhances SC activation and proliferation, favoring symmetric division and promoting differentiation at the expense of self-renewal. Therefore fibroblast-derived niche cues, including ECM deposition and soluble factors, critically influence SC fate and likely contribute to age-associated shifts in SC division dynamics. Given the pronounced effects of fibroblasts from aged mice on SC behavior regardless of myofiber donor age, we asked which fibroblast-derived niche factors drive the aging-associated changes in SC fate.

### 2.7. Fibronectin deposition by fibroblasts from aged mice drives loss of SC quiescence and promotes activation and differentiation

We asked if fibronectin deposition by fibroblasts from aged mice drives loss of SC quiescence and induces differentiation by altering fibronectin expression in heterochronic co-encapsulated 3D cultures. Myofibers from young or aged mice were co-encapsulated in 1 mM RGD hydrogels with fibroblast isolated from i) young mice and treated *in vitro* with scrambled siRNA control, ii) aged mice and treated *in vitro* with scrambled siRNA control, or iii) aged mice and treated *in vitro* with *Fn1* siRNA. Constructs were cultured for 2 days, with a 2-hour EdU pulse administered prior to fixation to assess SC proliferation and cell state based on Pax7 (stemness) and MyoD (activation) immunoreactivity **(Figure 7A and 7B)**.

**Figure 7.**
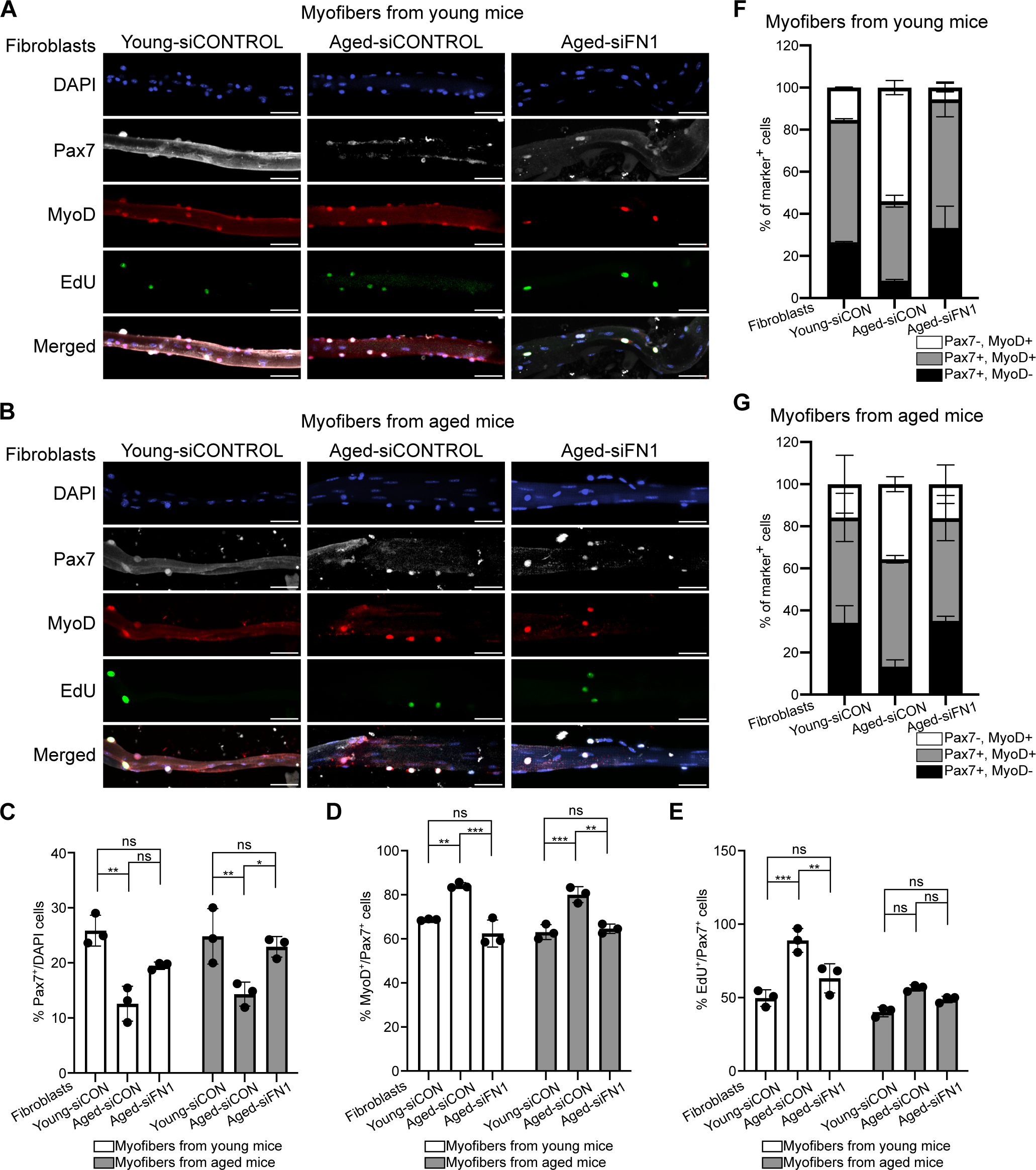
*Fn1* knockdown in fibroblasts from aged mice partially restores SC quiescence and suppresses differentiation. (A and B) Representative images of myofibers from young (A) or aged (B) mice co-encapsulated in 1 mM RGD hydrogels with fibroblasts from young mice treated with scrambled siRNA control, fibroblasts from aged mice treated with scrambled siRNA control, or fibroblasts from aged mice treated with *Fn1* siRNA. SCs identified by Pax7 and MyoD immunoreactivity, and assessed for EdU incorporation after 2 days in culture. Scale bars, 50 μm. (C-E) Quantification of Pax7^+^ (C), MyoD^+^ (D), and EdU^+^ (E) SCs on myofibers from young or aged mice under the indicated conditions. (F and G) Distribution of SC states, defined as Pax7+/MyoD– (quiescent), Pax7+/MyoD+ (myoblasts), and Pax7–/MyoD+ (differentiated myoblasts), on myofibers from young (F) or aged (G) mice under the indicated conditions. For all quantifications, *n* > 9 myofibers analyzed with samples from 3 mice, and statistics performed based on 3 independent mice; data are presented as mean ± SD; *\*P* < 0.05, *\*\*P* < 0.01, and *\*\*\*P* < 0.001 in a two-way ANOVA test.

SCs on myofibers from young mice co-encapsulated with fibroblasts from aged mice treated with scrambled siRNA control reduced Pax7⁺ cells (13 ± 3%) compared to co-encapsulation with fibroblasts from young mice treated with scrambled siRNA control (26 ± 3%), consistent with our previous data **(Figure 7C)**. The reduced number of Pax7^+^ cells was partially rescued by *Fn1* knockdown in fibroblasts from aged mice (19 ± 1%) but did not reach statistical significance. A similar trend was observed in SCs on myofibers from aged mice, where fibroblasts from aged mice reduced the proportion of Pax7⁺ cells (14 ± 2%) with *Fn1* knockdown restoring the Pax7⁺ SC population (23 ± 2%). Thus, fibroblasts in aged mice secrete a fibronectin-rich matrix that compromises SC maintenance and likely contributes to SC loss.

We evaluated SC activation and proliferation following siRNA-mediated *Fn1* knockdown to determine whether fibronectin mediates the fibroblast-induced shift to activate SCs. Control-treated fibroblasts from aged mice increased the fraction of Pax7^+^MyoD^+^ cells compared to control-treated fibroblasts from young mice regardless of the myofiber donor age **(Figure 7D)**. Knockdown of *Fn1* in fibroblasts from aged mice significantly attenuated this effect, restoring SC activation to levels similar to those in cultures with fibroblasts from young mice. A corresponding rescue of SC proliferation occurred on myofibers from young mice **(Figure 7E)**, where *Fn1* knockdown reduced the elevated EdU incorporation to levels similar to those with fibroblasts from young mice. In contrast, SC proliferation on myofibers from aged mice remained relatively constant, with no significant differences across fibroblast conditions. Fibronectin-rich fibroblasts from aged mice promote SC activation and enhance proliferation, shifting SCs toward an activated state in myofibers from young mice.

Co-encapsulation of fibroblasts from aged mice treated with scrambled siRNA control altered SC fate on young myofibers, depleting the quiescent pool and biasing cells toward differentiation **(Figure 7F)**. Notably, *Fn1* knockdown in fibroblasts from aged mice fully reversed this age-driven depletion of quiescence, restoring both quiescent (Pax7+/MyoD–; 33 ± 10%) and differentiated (Pax7–/MyoD+; 6 ± 2%) populations to baseline levels observed with fibroblasts from young mice. The fibronectin-dependent rescue was equally effective on myofibers from aged mice **(Figure 7G)**, where silencing *Fn1* again returned quiescent and differentiated SC proportions to those observed in cultures with fibroblasts from young mice (35 ± 2% and 16 ± 9%, respectively). Therefore, fibronectin produced by fibroblasts from aged mice is a primary driver disrupting SC cell state balance; depleting fibronectin in fibroblasts from aged mice is sufficient to rescue SC maintenance regardless of the age of the myofiber donor.

We identify fibroblast-derived fibronectin as a key ECM regulator of SC state, linking age-associated matrix remodeling to impaired SC maintenance. Using a defined 3D co-encapsulation system, we demonstrate that fibroblasts from aged mice shift the SC population away from quiescence and toward terminal differentiation. Given that quiescence is essential for long-term stem cell self-renewal,^[6, 39]^ the shift of SCs toward differentiation likely contributes to depletion of the SC pool. These effects are substantially reversed by *Fn1* knockdown in fibroblasts from aged mice, establishing excessive fibronectin deposition as a critical ECM-level driver of SC dysfunction during aging. Excessive fibronectin accumulation likely alters integrin-mediated adhesion and mechanotransduction,^[5, 40]^ thereby reshaping SC signaling thresholds that govern the balance between self-renewal and differentiation. Collectively, these changes bias SC fate decisions toward premature SC commitment and regenerative exhaustion at the expense of quiescence maintenance.

Loss of regenerative responses in skeletal muscle is not solely SC-intrinsic but is actively reinforced by fibroblasts that establish a pro-differentiation, quiescence-depleting niche by accumulating fibronectin. Our conclusions align with prior data where transplanting SCs from young mice into aged mouse muscle failed to rescue age-associated self-renewal defects.^[6]^ In addition, SCs from young mice expand poorly on fibrotic, collagen-rich ECM derived from aged mouse muscle.^[23]^ Therapeutically, fibronectin accumulation and fibronectin-integrin signaling are thus potential targets for restoring SC homeostasis in aged muscle; however, given the broad expression of fibronectin across tissues, effective interventions will likely require localized delivery strategies. Intramuscular delivery of *Fn1-*targeting siRNAs may be a potential strategy to limit fibronectin accumulation, thereby preventing premature SC differentiation and maintaining a functional and responsive SC pool during aging.

We assessed the effects of fibroblasts on SCs using a novel material-based co-culture system that could be extended to other muscle-resident cell populations to investigate their relative contributions to SC behavior in regenerating muscle in young and aged organisms. For example, incorporation of immune cells, endothelial cells, or pericytes would allow one to further dissect multicellular signaling networks that shape the SC niche and direct SC behavior. Beyond mechanistic studies, our *in vitro* platform might provide a more physiologically relevant system to screen therapeutic targets to inhibit pathological ECM remodeling or restore age-associated SC niche dysfunction.

## 3. Conclusion

Fibronectin remodeling is transient in regenerating muscles of young mice but is prolonged and dysregulated in muscles of aged mice. The altered ECM composition observed in aged mouse muscle is partly driven by intrinsic changes in fibroblasts, as aged mouse muscle contains increased numbers of fibroblasts and a higher proportion of activated fibroblasts *in vivo*. In fibroblasts from aged mice, ⍺SMA expression is elevated and fibronectin deposition is enhanced *in vitro*. We established an *in vitro* co-encapsulation system using a fully defined synthetic viscoelastic hydrogel preserving native SC polarity and the SC-myofiber-matrix interface to determine whether aging affects SC fate via fibroblast interactions. Using our novel co-culture platform, fibroblasts from aged mice remain intrinsically more activated than those from young mice, independent of the age of the myofiber donor. Fibroblasts from young mice support SC self-renewal, whereas fibroblasts from aged mice bias SCs toward symmetric division and differentiation. Notably, *Fn1* knockdown in fibroblasts from aged mice partially restores SC quiescence and reduces activation and differentiation. Excessive fibronectin deposition by fibroblasts in aged mouse muscle is a key driver of niche dysfunction that disrupts ECM remodeling, alters SC fate, and ultimately impairs regenerative capacity, highlighting fibroblast-derived ECM as a targetable axis for rejuvenating muscle repair.

## 4. Experimental Section/Methods

### Animals

All mice were bred and housed according to National Institutes of Health (NIH) guidelines for the ethical treatment of animals in a pathogen-free facility at the University of Colorado at Boulder. All animal protocols and procedures were approved by the University of Colorado Institutional Animal Care and Use Committee, and the conducted studies complied with all ethical regulations. Wild-type mice were C57BL/6 (Jackson Labs, ME, USA). Mice from 4-6 months of age were considered young, and mice from 20-24 months of age were considered aged.

### Animal procedures

Mice were anesthetized with isoflurane, followed by injection of 50 μL 1.2% BaCl₂ into the left tibialis anterior (TA) muscle, and then injured TA muscles were harvested 4 and 7 days post-injury.

### Immunohistochemistry of tissues

TA muscles (uninjured, 4 dpi, and 7 dpi) of young and aged mice (C57BL/6J) were dissected and fixed with 4% paraformaldehyde (PFA) for 2 hours on ice. Muscles were then rinsed with phosphate buffered saline (PBS) and transferred to PBS with 30% sucrose at 4 °C overnight. Muscle tissues were embedded in optimal cutting temperature (O.C.T.) compound on dry ice and cryo-sectioned using a Leica cryostat to generate 15-μm-thick cross-sections. After rinsing with PBS, sections were fixed with 4% PFA at room temperature for 8 minutes. Heat-induced epitope retrieval was performed by immersing post-fixed slides in 10 mM citrate buffer with 0.05% Tween-20 (pH 6.0) and heating them under high pressure for 6 minutes using a Cuisinart model CPC-600 pressure cooker. Then sections were washed with PBS and permeabilized with 0.25% Triton X-100 and 2% bovine serum albumin (BSA) in PBS for 1 hour at room temperature. Primary antibodies included rabbit anti-fibronectin (Abcam), mouse anti-Pax7 (Developmental Studies Hybridoma Bank, University of Iowa, USA), mouse anti-⍺-SMA (Abcam), and goat anti-PDGFRA (R&D Systems) all used at a 1:250 dilution in PBS containing 2% BSA. Alexa Fluor secondary antibodies (Thermo Fisher Scientific) were used at a 1:1000 dilution in PBS containing 2% BSA, with 1 μg/mL 4′,6-diamidino-2-phenylindole (DAPI) added. Samples were mounted with Mowiol supplemented with DABCO (Sigma-Aldrich) as an anti-fade agent.

### Software packages

- CellRanger Software Suite/3.0.1
- FastQC 0.11.8
- R 4.3.3
- Seurat 5.1.0
- CellChat 1.6.1
- SoupX 1.5.2

### Quality control, read alignment, and expression quantification

*FastQC* was used to evaluate the quality of read depth on each replicate of Fastq files from sequencing. *CellRanger* was employed to process Fastq files by aggregating technical replicates and generating gene count matrices. Transcriptomes from snRNA-seq were aligned using a previously validated custom pre-mRNA mm10 reference package.

### Accounting for experimental noise and doublets

*CellRanger*-aligned feature matrices were loaded into *R* using the load10X() function from the *SoupX* package. AutoEstCont() was used to predict RNA contamination and adjustcounts() normalized counts to an estimated noise parameter. Manual correction of gene expression matrices removed doublets and debris using recommended metrics. In addition, cells or nuclei expressing more than 2,500 or less than 200 unique features were removed.

### Normalization, dimensional reduction, and nuclear clustering

The tutorial provided by the Mayaan lab (satijalab.org/seurat/articles/pbmc3k_tutorial) was followed to normalize data and mitochondria and low-quality nuclei were removed. Seurat objects were processed using the NormalizeData(), FindVariableFeatures(), and ScaleData() functions from *Seurat* to scale, and log normalize gene counts within each sample. Then, the *Seurat* functions FindIntegrationAnchors() and IntegrateData() were used to integrate data using reciprocal PCA (rPCA). Nuclear clusters were separated using the shared nearest neighbor (SNN) modularity optimization-based clustering. The minimum number of neighbors, minimum distance between neighbors, and resolution parameters were adjusted while performing dimensional reduction of the integrated *Seurat* object using UMAP.

### CellChat analysis of cell-cell communication

Cell-cell communication analysis of each *Seurat* object was performed using *CellChat* (v1.6.1) following the workflow.^[41]^ ComputeCommunProb() was used with an 18% Truncated Mean cutoff to determine the cell-type specific expression of ligand or receptors in a pathway of interest.

### Fibroblast isolation and culture

To isolate fibroblasts, hindlimb muscles were dissected, minced into a fine paste, and digested in Ham’s F-12 (Gibco) containing 400 U/mL collagenase (Worthington) at 37 °C for 1 hour with vigorous shaking every 10 minutes. Collagenase was then quenched using Ham’s F-12 supplemented with 15% horse serum (Gibco), 1% penicillin-streptomycin (Gibco), and 0.8 mM CaCl_2_. The muscle digest was sequentially filtered through 100-, 70-, and 40-μm strainers (Thermo Fisher Scientific), centrifuged at 200 × g for 5 minutes, and resuspended in Ham’s F-12 containing 15% horse serum, 1% penicillin-streptomycin, and 0.8 mM CaCl_2_. Cells were pre-plated onto untreated petri dishes for 1 hour to allow fibroblast attachment. After pre-plating, non-adherent cells were removed, and adherent fibroblasts were cultured in fresh Ham’s F-12 containing 15% horse serum, 1% penicillin-streptomycin, 0.8 mM CaCl_2_, and 0.5 nM fibroblast growth factor 2 at 5% CO_2_ and 37 °C.

### siRNA knockdown of Fn1

Fibroblasts were harvested by 5 min of trypsinization and seeded at 2 ⨉ 10^5^ cells per well in 6-well tissue culture plates. After 24 h, cells were washed once with siRNA transfection medium (Santa Cruz Biotechnology). Fibronectin siRNA (60 pmol, Santa Cruz Biotechnology) or control siRNA (60 pmol, Santa Cruz Biotechnology) was transfected using siRNA transfection reagent and medium (Santa Cruz Biotechnology) according to the manufacturer’s instructions and incubated at 37 °C with 5% CO_2_ for 6 h. Following transfection, the medium was replaced with Ham’s F-12 supplemented with 15% horse serum, 1% penicillin-streptomycin, 0.8 mM CaCl_2_, and 0.5 nM fibroblast growth factor 2. Cells were allowed to recover overnight and used for co-encapsulation the following day.

### Myofiber isolation

Extensor digitorum longus muscles were dissected and digested in Ham’s F-12 (Gibco) containing collagenase (400 U/mL) (Worthington) at 37 °C for 1.5 hours with gentle rotation. After digestion, collagenase was inactivated with Ham’s F-12 containing 15% horse serum (Gibco) and 0.8 mM CaCl_2_. Then, a flame-polished glass pipet was used to transfer individual myofibers to 6-well tissue culture plates containing Ham’s F-12 supplemented with 15% horse serum, 1% penicillin-streptomycin (Gibco), 0.8 mM CaCl_2_, and 0.5 nM fibroblast growth factor 2. Myofibers were cultured in suspension at 37 °C and 5% CO₂ for 24 hours prior to encapsulation.

### Synthesis of HA-hydrazide and characterization

HA-hydrazide was synthesized as previously described.^[38, 42]^ Briefly, HA (60 kDa, 500 mg) was dissolved in dH₂O (100 mL), followed by the addition of adipic acid dihydrazide (≈6.5 mg, >60-fold molar excess) while adjusting the pH to 6.8. 1-Ethyl-3-(3-dimethylaminopropyl)carbodiimide (EDC, 776 mg) and hydroxybenzotriazole (HOBt, 765 mg), separately dissolved in a 1:1 DMSO/dH₂O mixture, were added dropwise to the reaction. The pH was maintained at 6.8 every 30 min for 4 h, and the reaction proceeded for 24 h. The product was dialyzed against dH₂O (8000 MWCO) for 3 days and lyophilized for 3 days. Then the product was redissolved in 5 wt.% NaCl/dH₂O, precipitated in ethanol, redialyzed for three days, and lyophilized for an additional three days. The final white powder (430 mg, 86% yield) was flash frozen in liquid nitrogen and stored at −20 °C. Molecular weight (∼86 kDa) was confirmed by GPC characterization and functionalization (∼28.5%) was confirmed by ¹H NMR.^[15]^

### Synthesis of HA-aldehyde and characterization

HA-aldehyde was synthesized as previously described.^[38, 42]^ Briefly, HA (500 kDa, 500 mg) was dissolved in dH_2_O (50 mL), and then sodium periodate (267.5 mg) was added into the reaction mixture. After stirring in the dark for 2 h, the reaction was quenched with ethylene glycol (70 µL), dialyzed against dH₂O (8000 MWCO) for 3 days, and lyophilized for another 3 days to obtain a white powder (471 mg, 94% yield). HA-aldehyde was flash frozen in liquid nitrogen and stored at −20 °C. The functionalization of the HA-aldehyde macromer was quantified using a 2,4,6-trinitrobenzene sulfonic acid (TNBS) assay as previously describe.^[43, 44]^ Briefly, the HA-aldehyde was dissolved at 2 wt.% (w/v) in dH₂O and reacted with tert-butyl carbazate (t-BC, in 1% trichloroacetic acid). After 24 h, the HA-aldehyde/t-BC and t-BC standards were reacted with 0.5 mL of TNBS solution (6 mM in 0.1 M sodium tetraborate, pH 8) for 1 h. The reaction was then quenched with 0.5 N hydrochloric acid. Molecular weight (∼148 kDa) was confirmed by GPC characterization and functionalization (∼36%) was confirmed by absorbance measured at 340 nm using a microplate reader.^[15]^

### PEG-bicyclononyne synthesis

PEG-bicyclononyne (BCN) was synthesized as previously described.^[38, 45]^ Briefly, 8-arm PEG-amine (40 kDa, 1.0 g, 0.2 mmol amine) and BCN-oSu (0.1 g, 0.343 mmol) were added to a 50 mL round-bottom flask. The components were dissolved in anhydrous DMF (10 mL), and the solution was stirred at room temperature under argon. N,N-Diisopropylethyleamine (0.8 mmol) was added, and the reaction proceeded overnight. The mixture was then diluted with dH_2_O and dialyzed for 3 days (8000 MWCO). The product was lyophilized for 3 days to yield a white powder (0.985 g, 98% yield). End-group functionalization (>95%) was confirmed by ¹H NMR.^[15]^

### Hydrogel Formulation

Hydrogels were prepared by dissolving functionalized HAs in PBS at 5 wt.%. Functionalized PEG-BCN was dissolved in PBS at 10 wt.%, and the benzaldehyde-PEG_3_-azide (BroadPharm) crosslinker was prepared at 20 mM in PBS. Final hydrogels were formed with a polymer content of 5 wt.%, based on stoichiometric calculations. In the reported formulation, 12% of the hydrazide groups on HA were substituted with benzaldehyde-PEG_3_-azide, yielding a hydrogel composed of 88% adaptable alkyl-hydrazone crosslinks and 12% irreversible azide-alkyne crosslinks, with PEG-BCN and HA-aldehyde incorporated at corresponding molar ratios.

### Myofiber and fibroblast co-encapsulation

The hydrogels were formed by pre-reacting the 5 wt.% HA-hydrazide with the benzaldehyde-PEG_3_-azide (20 mM) and the benzaldehyde-KRGDS (1 mM) in PBS overnight at 4 °C. The HA-aldehyde, PEG-BCN and PBS solutions were mixed. Fibroblasts were collected by a 5-min trypsinization and resuspended in the HA-hydrazide mixture at a density adjusted to yield 10,000 cells per 30 μL hydrogel upon addition of HA-aldehyde. Myofibers were picked using a flame-polished glass pipet and placed into a glass-bottom 96-well plate (Cellvis), with approximately 15 myofibers per well. After allowing the myofibers to settle in the incubator for 10 minutes, excess medium was carefully removed. Immediately afterward, the HA-hydrazide mixture containing fibroblasts was added to the well, followed by addition of the HA-aldehyde mixture to initiate gelation around the myofibers and fibroblasts. Myofibers were gently resuspended in 30 μL of hydrogel solution. After polymerization for 20 minutes in the incubator, fresh Ham’s F-12 supplemented with 15% horse serum, 1% penicillin-streptomycin, 0.8 mM CaCl_2_, and 0.5 nM fibroblast growth factor 2 was added. Cultures were maintained at 37 °C in an incubator with 5% CO_2_.

### Proliferation assays and immunocytochemistry

For proliferation assays, hydrogels containing myofibers and fibroblasts were treated with 10 μM EdU (Thermo Fisher Scientific) for 2 hours after two days of culture. Samples were then briefly washed with PBS and fixed with 10% formalin for 30 minutes. Following PBS washes, samples were permeabilized with 0.5% Triton X-100 and 4.6 mM N_3_-PEG_3_-OH in PBS for two hours, then incubated with the Click-iT reaction cocktail from the Click-iT EdU Alexa Fluor 488 kit (Thermo Fisher Scientific) for 30 minutes. After PBS washes, samples were incubated with 20 mM NH_4_Cl in PBS for 1 hour and blocked with 5% BSA in PBS at 4 °C overnight. For samples not subjected to EdU labeling, the same fixation, permeabilization, NH_4_Cl quenching, and blocking steps were performed, omitting the EdU incubation and Click-iT labeling. Primary antibodies included Pax7 (mouse, Developmental Studies Hybridoma Bank), MyoD (rabbit, Santa Cruz Biotechnology), fibronectin (rabbit, Abcam), and ⍺-SMA (mouse, Abcam), all used at a 1:250 dilution in PBS containing 5% BSA. Alexa Fluor secondary antibodies (Thermo Fisher Scientific) were used at a 1:500 dilution in PBS containing 5% BSA, with 1 μg/mL 4′,6-diamidino-2-phenylindole (DAPI) added. Phalloidin was added together with the secondary antibodies at a 1:500 dilution.

### Imaging and image analysis

Images of immunostained samples were carried out using a Nikon A1R Confocal Microscope equipped with a x20 NA = 0.75 objective. Images were visualized and analyzed with ImageJ. The brightness and contrast of each channel was separately adjusted for better visualization.

### Statistical analysis

Statistics were analyzed using Prism (GraphPad), and *P* < 0.05 was considered significant. For tibialis anterior muscle sections, at least 9 images were analyzed from 3 independent mice, and two-way ANOVA tests were performed based on the three biological replicates. For protein expression of fibroblasts on gelatin-coated plates, at least 150 cells were analyzed from 3 independent mice, and two-tailed unpaired student’s *t*-tests were performed based on the three biological replicates. For protein expression of fibroblasts co-encapsulated with myofibers in hydrogels, at least 9 images were analyzed from 3 independent mice, and two-way ANOVA tests were performed based on the three biological replicates. For hydrogel-encapsulated myofibers, at least 9 myofibers were analyzed from 3 independent mice, and two-way ANOVA tests were performed based on the three biological replicates.

## Acknowledgements

This work was supported by grants from the NIH (DE016523 and DK120921) to KSA and NIH (AR049446 and AR070630) to BBO. The imaging work was performed at the BioFrontiers Institute’s Advanced Light Microscopy Core (RRID: SCR_018302). Laser scanning confocal microscopy was performed on a Nikon A1R microscope supported by NIST-CU Cooperative Agreement award number 70NANB15H226.

## Data Availability Statement

This paper does not report original code. All data needed to evaluate the conclusions in the paper are present in the paper and/or the Supplementary Materials. Any additional information required to reanalyze the data reported in this paper are available from the corresponding author upon reasonable request

